# Morphometric Similarity Networks Detect Microscale Cortical Organisation And Predict Inter-Individual Cognitive Variation

**DOI:** 10.1101/135855

**Authors:** Jakob Seidlitz, František Váša, Maxwell Shinn, Rafael Romero-Garcia, Kirstie J. Whitaker, Petra E. Vértes, Paul Kirkpatrick Reardon, Liv Clasen, Adam Messinger, David A. Leopold, Peter Fonagy, Raymond J. Dolan, Peter B. Jones, Ian M. Goodyer, the NSPN Consortium, Armin Raznahan, Edward T. Bullmore

## Abstract

Macroscopic cortical networks are important for cognitive function, but it remains challenging to construct anatomically plausible individual structural connectomes from human neuroimaging. We introduce a new technique for cortical network mapping, based on inter-regional similarity of multiple morphometric parameters measured using multimodal MRI. In three cohorts (two human, one macaque), we find that the resulting morphometric similarity networks (MSNs) have a complex topological organisation comprising modules and high-degree hubs. Human MSN modules recapitulate known cortical cytoarchitectonic divisions, and greater inter-regional morphometric similarity was associated with stronger inter-regional co-expression of genes enriched for neuronal terms. Comparing macaque MSNs to tract-tracing data confirmed that morphometric similarity was related to axonal connectivity. Finally, variation in the degree of human MSN nodes accounted for about 40% of between-subject variability in IQ. Morphometric similarity mapping provides a novel, robust and biologically plausible approach to understanding how human cortical networks underpin individual differences in psychological functions.

## Introduction

Despite decades of neuroscience research using magnetic resonance imaging (MRI), there is still a lack of validated and widely-accessible tools for mapping the large-scale network architecture of anatomically connected regions in an individual human brain. There are currently two standard approaches available for imaging anatomical connectivity in humans: tractography from diffusion-weighted imaging (DWI) and structural covariance network (SCN) analysis.

Diffusion-weighted tractography seeks to reconstruct the trajectory of axonal tracts from the principal directions of the diffusion of water molecules, which tend to move in parallel to bundles of nerve fibres. This technique applies to data collected from a single participant and is a powerful tool for elucidating localised patterns of anatomical connectivity. However, it remains challenging to use tractography to map connectivity between *all* brain regions because long distance projections (e.g., between bilaterally homologous areas of cortex via the corpus callosum) are systematically under-recovered (Dauguet et al., 2007; Donahue et al., 2016). Moreover, there is growing concern that the statistical analysis of diffusion-weighted data is compromised by head movement (Walker et al., 2012) and by a large number of false positive connections (Maier-Hein et al., 2016; Thomas et al., 2014). Future improvements seem likely to depend, in part, on advances in scanner design and image acquisition methods, which may become increasingly the domain of a few highly-specialised centres (Lerch et al., 2017).

Structural covariance analysis uses simpler measurements to reconstruct whole brain networks; but the neurobiological interpretation of structural covariance networks (SCN) is problematic and, crucially, this method typically depends on MRI data collected from a large number of participants. The basic idea of structural covariance analysis is simple: a single morphometric feature, like cortical thickness, is measured at each region in multiple images. Then the covariance (usually, in fact, the correlation) between regional estimates of cortical thickness is estimated for each possible pair of regions, resulting in a single SCN for the whole group (Alexander-Bloch et al., 2013a). Despite the existence of methods for generating SCNs in individual subjects (Batalle et al., 2013; Kong et al., 2015; Li et al., 2017; Tijms et al., 2012), these techniques have been restricted to the use of morphometric variables available through standard structural T1-weighted (T1w) MRI sequences.

Here, we explore a different approach to human cortical network mapping which leverages the growing capacity to extract multiple different anatomical indices across multiple imaging modalities (Lerch et al., 2017). Rather than estimating the inter-regional correlation of a single macro-structural variable (like cortical thickness or volume) measured in multiple individuals (structural covariance analysis), we estimated the inter-regional correlation of multiple, macro- and micro-structural multimodal MRI variables in a single individual (morphometric similarity mapping). This novel strategy integrates three complementary strands of research for the first time.

First, there is histological evidence from non-human primates that axo-synaptic connectivity is stronger between micro-structurally similar cortical regions than between cytoarchitectonically distinct areas (Barbas, 2015; Goulas et al., 2017; Goulas et al., 2016). Second, there is encouraging evidence that conventional MRI sequences can serve as proxy markers of cortical microstructure. Cortical MRI metrics – such as magnetization transfer (MT), a marker of myelination - show spatial gradients in humans (Glasser et al., 2016) which align closely with known histological gradients in non-human primates (Wagstyl et al., 2015). Third, there is emerging evidence that structural properties of the human cortex are more precisely estimated by the combined analysis of more than one MRI morphometric index at each region (e.g. cortical thickness and sulcal depth (Vandekar et al., 2016), cortical thickness and myelination (Glasser and Van Essen, 2011; Whitaker et al., 2016), or cortical thickness and grey matter volume (Sabuncu et al., 2016)). On this basis, we predicted that morphometric similarity mapping with multiple MRI morphometric indices could provide a new way of estimating the linked patterns of inter-regional histological similarity and anatomical connectivity within an individual human brain.

First we demonstrated the feasibility of morphometric similarity mapping and network analysis of multi-parameter MRI data, by estimating individual human brain structural network properties for each member of a cohort of healthy young people (N=296). Then we assessed the robustness of the methods and results to variation in data acquisition and pre-processing parameters; and assessed replicability by analysis of a second, independent human MRI dataset (N=124). To test the biological validity of human MRI-based morphometric similarity mapping, we focused on two hypotheses: i) that the edges between nodes in each morphometric similarity network (MSN) linked cortical areas of the same cytoarchitectonic class; and ii) that MSN edges linked nodes with high levels of co-expression of genes specialised for neuronal functions. We used MRI and tract-tracing data of the macaque cortex to demonstrate the generalisability of the methods to non-human species and to test a third key biological hypothesis: (iii) that MSN edges link nodes that are directly, anatomically connected by an axonal projection from one cortical area to another. Finally, on the basis of these critical foundational steps, we tested a fourth hypothesis: iv) that inter-individual differences in human morphometric similarity networks are related to differences in intelligence (IQ).

## Results

### Morphometric similarity matrices

To investigate the feasibility of morphometric similarity mapping, we first analysed MRI data on 10 morphometric variables measured at each of 308 cortical regions in a primary cohort of 296 healthy young adults. These data were collected as part of the Neuroscience in Psychiatry Network (NSPN) collaboration between the University of Cambridge and University College London (see **Methods**).

The morphometric similarity analysis pipeline (**Figure 1**) transformed each individual’s set of multimodal MRI feature maps into a morphometric similarity matrix of pair-wise inter-regional Pearson correlations of morphometric feature vectors. The strength of association (morphometric similarity) between regions decayed exponentially as a function of increasing anatomical (Euclidean) distance between areas (median R^2^_*Adjusted*_ across 296 subjects = 0.05, range = 0.02-0.13, σ = 0.02, all *P* < 0.001). This was also the case in the sample mean MSN (R^2^_*Adjusted*_ = 0.15, *P* < 0.001) (see **Figure S4A**).

**Figure 1.**
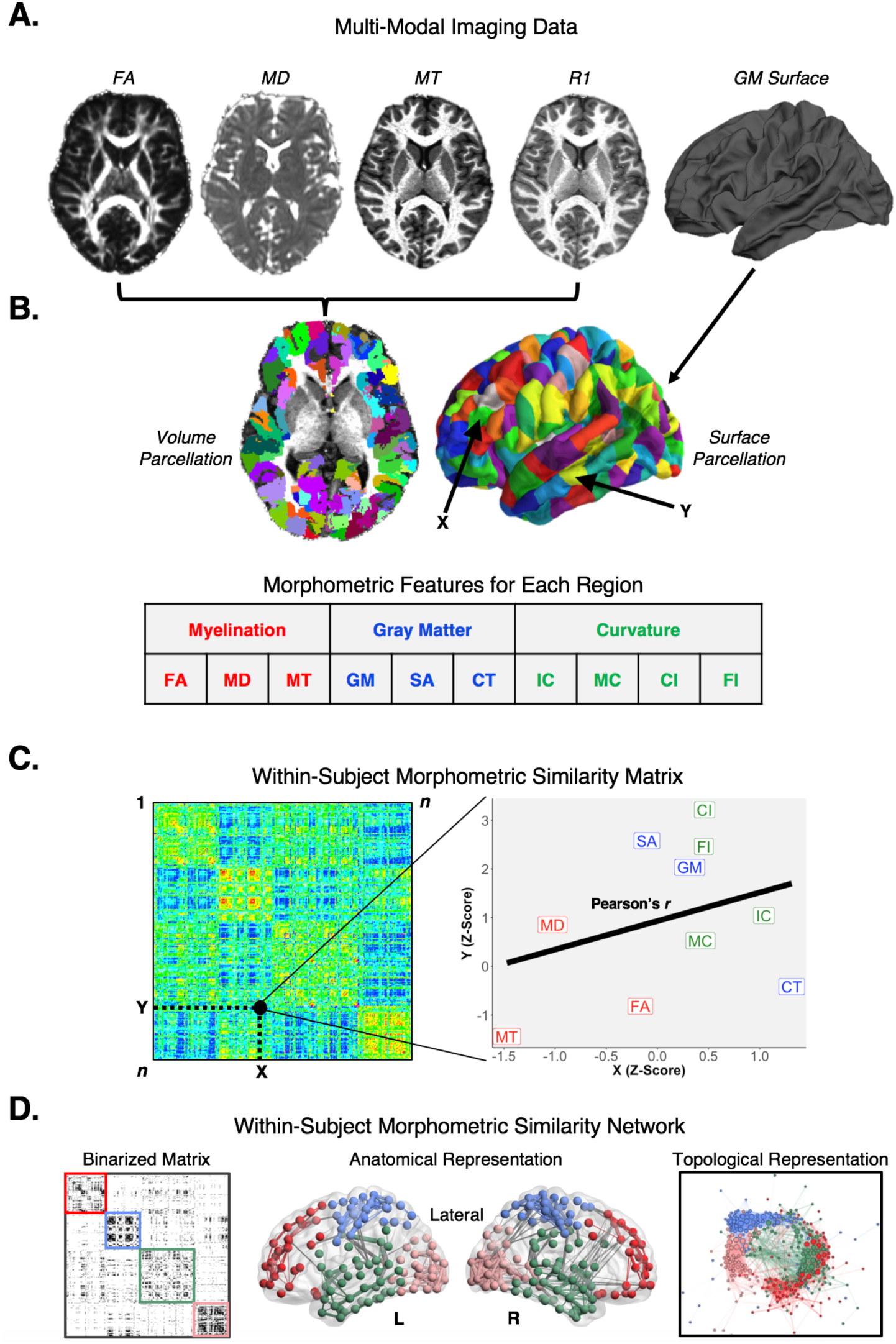
The morphometric similarity network processing pipeline. **A)** Multiple MRI parameters were available from MRI and DWI data on each subject. **B)** All MRI data were mapped to the same cortical parcellation template, which comprised 308 sub-regions of the Desikan-Killiany atlas with approximately equal surface area. 10 regional morphometric features were estimated and normalised to produce a 10 × 308 feature matrix for each subject. **C)** The morphometric similarity between each possible pair of regions was estimated by the Pearson’s correlation between their morphometric feature vectors to produce a 308 × 308 morphometric similarity matrix. **D)** Morphometric similarity networks are binary graphs constructed by thresholding the morphometric similarity matrix so that the strongest (supra-threshold) edges are set equal to 1 (and all others set equal to 0). The organisation of MSNs can be visualised (from left to right) in matrix format, in anatomical space, or in a topological representation where nodes are located close to each other if they are connected by an edge. FA = fractional anisotropy, MD = mean diffusivity, MT = magnetization transfer, GM = grey matter volume, SA = surface area, CT = cortical thickness, IC = intrinsic (Gaussian) curvature, MC = mean curvature, CI = curved index, FI = folding index.

For each individual, we also estimated the morphometric similarity of each region to the rest of the regions in the brain, simply by averaging the edge weights connecting it to all other nodes (i.e., the average of the off-diagonal elements of a row or column in the morphometric similarity matrix). Consistently across individuals, the regions with morphometric profiles more similar, on average, to all other regions (i.e., high nodal similarity values) were located in frontal and temporal association cortex; whereas regions with more distinctive morphometric profiles compared, on average, to other regions (i.e., low nodal similarity) were located in occipital cortex (**Figure 2A; Figure S1**). Nodal similarity in the sample mean MSN was also patterned anatomically as in typical individual MSNs (**Figure 2A; Figure S1**), with highest nodal similarity concentrated in frontal and temporal cortex (**Figure 2B**).

**Figure 2.**
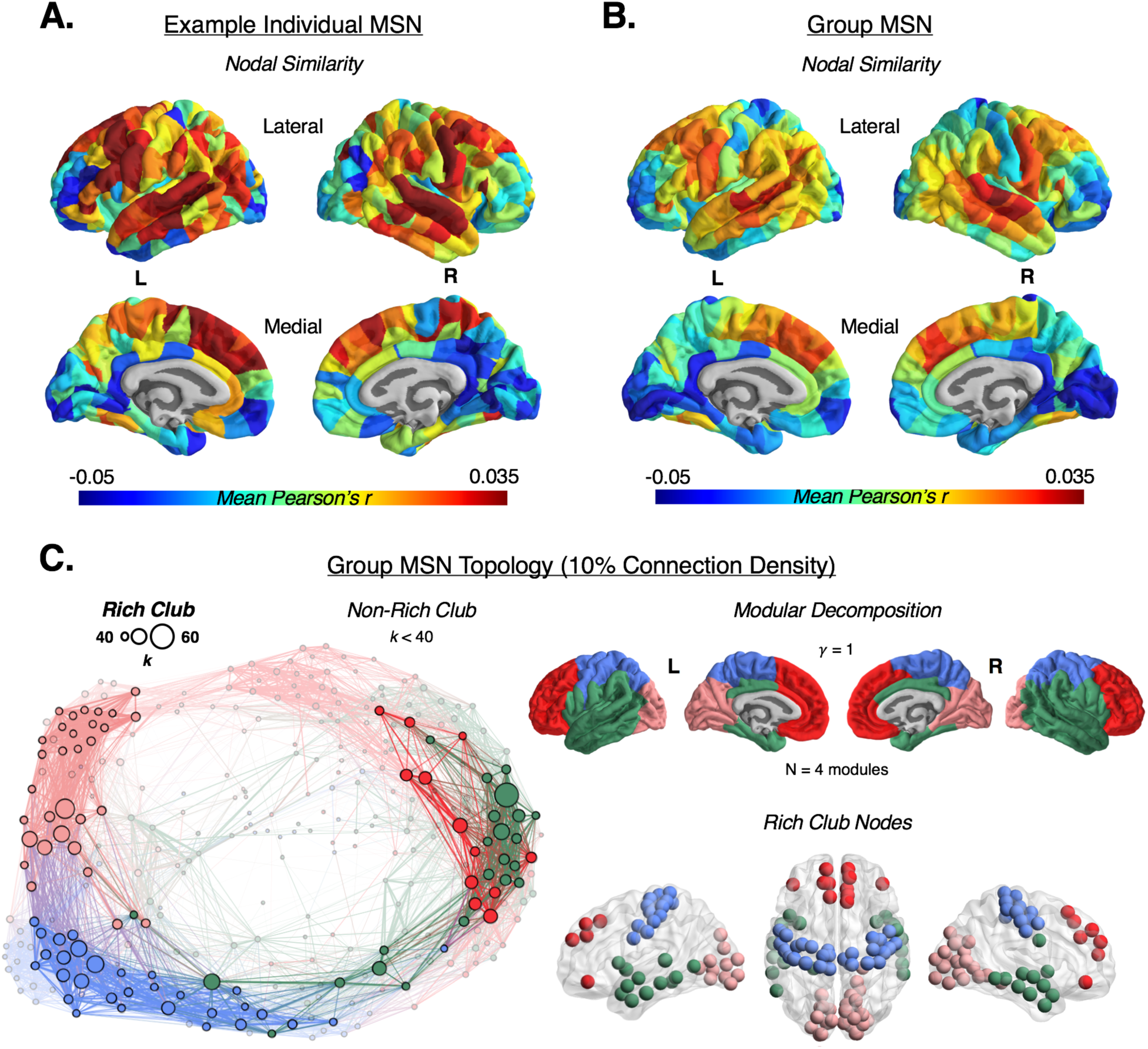
Morphometric similarity networks. Spatial patterning of an individual **(A)** and group average **(B)** morphometric similarity matrix. For the individual and group matrix, the row means are plotted on the cortical surface of the template brain, representing the average morphometric similarity (Pearson’s *r*) of each node. The colour scale represents the mean nodal similarity. **C)** (Right, top) Modular partitioning of the group average morphometric similarity network (MSN), thresholded at 10% connection density, using the Louvain modularity algorithm. The *γ* resolution parameter dictates the number of detected modules; *γ* = 1 yielded four distinct spatially contiguous modules, which approximately correspond to the lobes of the brain. (Left) Topological representation of the group MSN, thresholded at 10% connection density, highlighting the rich club of densely inter-connected hub nodes (opaque). The size of the nodes is scaled according to degree, and the thickness of the edges is scaled according to edge weight. (Right, bottom) The rich club nodes are shown in their anatomical location and coloured according to modular affiliation. See also Figures S1-3.

### Morphometric similarity networks

We thresholded the individual morphometric similarity matrices to generate binary graphs or morphometric similarity networks (MSNs). To characterise the topology of these MSNs, we calculated graph metrics at a range of connection densities (10-40% in 5% increments) generated by thresholding the morphometric similarity matrices to include varying percentages of the most strongly positive edge weights or pair-wise inter-regional correlations (**Figure S3A**). At all connection densities, the individual MSNs consistently demonstrated a repertoire of complex topological features shared by diverse naturally-occurring networks (Barabási, 2016; Fornito et al., 2016) including: a fat-tailed (i.e., right- or positive-skewed) degree distribution, implying the existence of hub nodes; small-worldness (near-random path length or global efficiency combined with greater-than-random clustering); and a community structure comprising hierarchical modules and a rich club (**Figure 2**).

We next resolved the modular community structure of MSNs by partitioning the sample mean MSN from coarse-to fine-grained scales defined by the resolution parameter (*γ*) of a consensus (1000 runs) Louvain modularity algorithm (Lancichinetti and Fortunato, 2012) (**Figure 2C; Figure S2C**). At all scales of the modular hierarchy, the community structure consists of bilaterally symmetric and spatially contiguous modules tiling the cortex in a pattern that respects macro-structural features of the cortical sheet (**Figure 2C**). For example, the 6-module solution subdivides the temporal module of the 4-module solution into anterior, middle, and superior (encompassing the insula) portions of temporal cortex (**Figure S2C**). These results are qualitatively consistent with a prior study of the hierarchical patterning of the heritability of cortical surface area (Chen et al., 2012), suggesting that the topological community structure of morphometric similarity could arise, at least in part, from genetic contributions to regional brain morphology.

We also demonstrated another aspect of network community structure – a core/periphery organisation comprising a core of highly interconnected hub nodes (a rich club). The rich club of the sample mean MSN included hubs that were distributed across all network modules (**Figure 2C**) as has been previously reported for DWI networks (van den Heuvel and Sporns, 2011). Individual MSNs typically demonstrated a similar community structure (modules and rich club) to that of the sample mean MSN (**Figure S2D**).

### Consistency, robustness and replicability of MSNs

First, we established that individual MSNs show moderate-high consistency with the group average MSN, both in terms of (i) the average correlation between individual MSN edge weights and the sample mean MSN edge weights (average *r* = 0.60; SD = 0.05; all *P* < 0.001), and (ii) the average correlation between individual MSN nodal similarities and the sample mean MSN nodal similarities (average *r* = 0.66; SD = 0.10; all *P* < 0.001) (see **Figure 2; Figure S1)**.

Second, we quantified the robustness of MSNs to methodological variations including (i) a reduction in the number of morphometric features available for analysis, i.e. using only 5 T1-weighted features rather than all 10 features potentially estimated in the NSPN cohort; and (ii) construction of MSNs using MRI data collected at lower magnetic field strength (1.5T) in a second, independent cohort and pre-processed using different segmentation and parcellation tools.

We re-analysed the NSPN cohort of 296 participants using a reduced set of 5 morphometric features that can be derived from any T1-weighted MRI scan: cortical thickness (CT), surface area (SA), grey matter volume (GM), mean curvature (MC), and intrinsic (Gaussian) curvature (IC). We found that the 5-feature MSNs were very similar to the 10-feature MSNs: for example, the sample mean edge weights and nodal similarities were strongly correlated between the 5-feature and 10-feature MSNs (*r* = 0.68, *P* < 0.001 and *r* = 0.91, *P* < 0.001, respectively). However, the standard deviation for edge weights and nodal similarity was greater in the 5-feature MSNs (0.506 and 0.028, respectively) than the 10-feature MSNs (0.346 and 0.016, respectively) (**Figure S4B**), indicating greater precision of MSN estimation based on a larger number of parameters or features per regional node.

To test replicability, we used identical methods to construct MSNs from T1-weighted MRI data collected at 1.5T field strength from an independent cohort of 124 healthy participants (National Institutes of Health (NIH) MRI Study of Normal Brain Development (Evans and Brain Development Cooperative, 2006; Giedd et al., 1999; Giedd et al., 2015). As shown in **Figure S2** and **Figure S3,** the 5-feature NIH MSNs had comparably complex topology to the 5- and 10-feature MSNs from the NSPN cohort (i.e., small-worldness, hubs, modularity and a rich club). The standard deviation of edge-wise and nodal similarity statistics in the NIH 5-feature MSNs (0.431 and 0.028, respectively) was approximately the same as the NSPN 5-feature MSNs but greater than the standard deviations in the 10-feature MSNs (**Figure S4B**).

### Morphometric similarity and cortical cytoarchitecture

To test the first biological hypothesis that regions connected by an edge in a morphometric similarity network are more likely to belong to the same cytoarchitectonic class, we compared the anatomical distribution of network edges to the histological classification of cortical areas (Solari and Stoner, 2011; Vertes et al., 2016; von Economo and Koskinas, 1925; Whitaker et al., 2016). Our adapted cytoarchitectonic parcellation of human cortex (Vertes et al., 2016; Whitaker et al., 2016) defines 7 spatially contiguous and bilaterally symmetric cortical classes that are microscopically differentiated by cortical lamination patterns (**Figure 3A**). At a nodal level of analysis, we explored the distribution of nodal similarity in the individual MSNs in relation to the cytoarchitectonic classification of each node. Across subjects, there were significant differences in mean nodal similarity between classes (repeated-measures ANOVA, *F*(1, 2077) = 796.5, *P* < 0.001). Cytoarchitectonic classes 1, 2 and 3 (corresponding to motor and association cortices) comprised cortical areas with higher nodal similarity and degree than cytoarchitectonic classes 4, 5 and 6 (corresponding to primary and secondary sensory cortical areas) (**Figure 3A**). These results confirm that MSN hubs are predominantly located in motor and association cortical areas.

**Figure 3.**
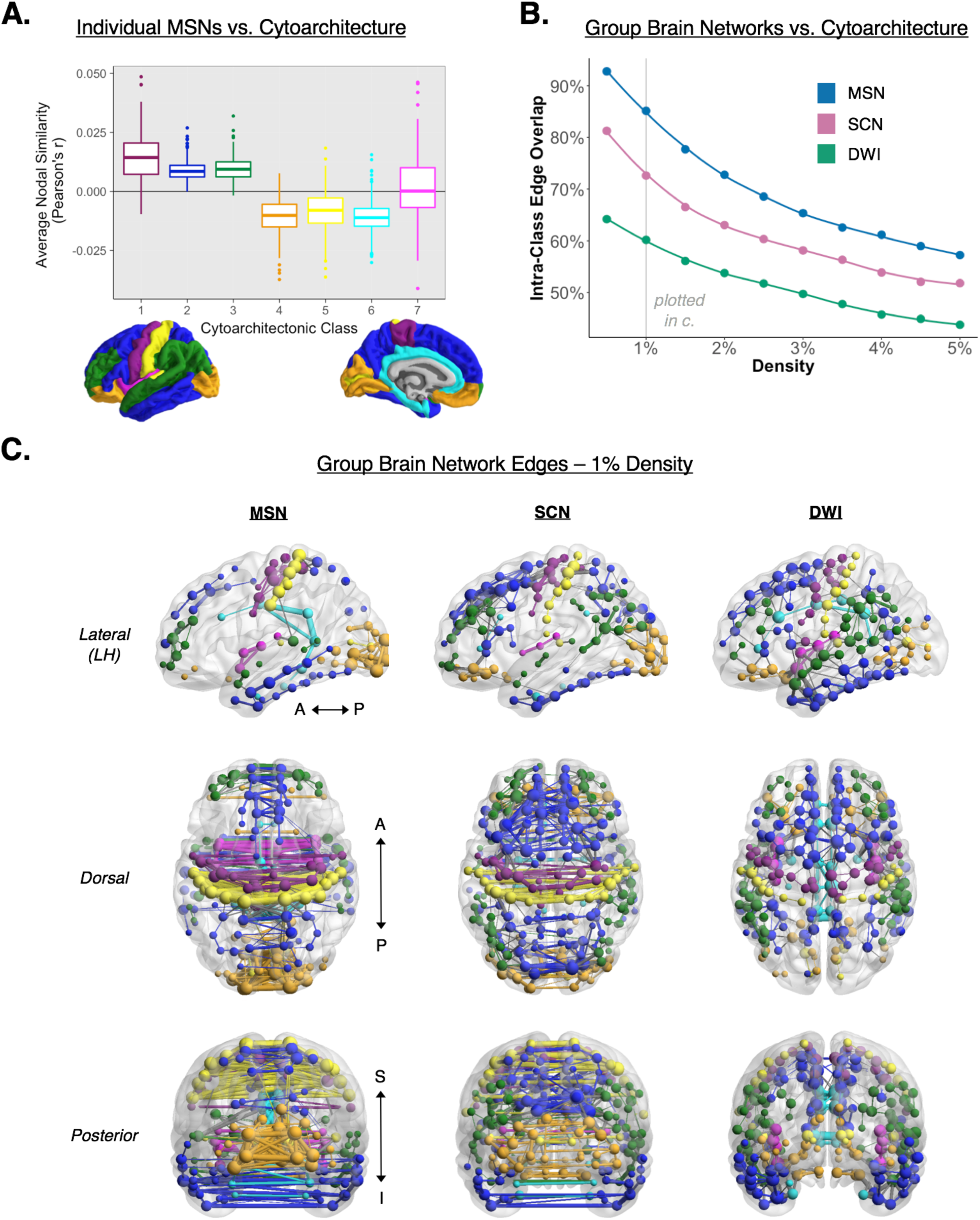
Comparison of morphometric similarity networks, and other MRI networks, to a cytoarchitectonic classification of cortex. **A)** Average nodal similarity scores for individual MSNs within each of the von Economo and Koskinas (1925) cortical classes plus the insula and cingulate cortex. Highest nodal similarity was consistently found in classes 1-3 (motor and association cortex) – areas with the most pyramidal neurons in supragranular layers of cortex. **B)** Proportion (percentage) of intra-class edges in the group MSN as a function of connection density (0.5-10%, 0.5% intervals). The MSN has a higher percentage of intra-class edges compared to the SCN and DWI networks at all densities, demonstrating the high correspondence of MSN topology with cortical cytoarchitectonics. **C)** Graphs of each of the MRI networks, thresholded at 1% density, with nodes and intra-class edges coloured according to cytoarchitectonic class and inter-class edges drawn in grey. The MSN shows the greatest connectivity between bilaterally symmetric cortical regions, relative to the SCN and DWI networks. Lower and upper bounds of the boxplots represent the 1^st^ (25%) and 3^rd^ (75%) quartiles, respectively. Left hemisphere = LH, Anterior = A, Posterior = P, Superior = S, Inferior = I.

The cytoarchitectonic parcellation of cortex also provided a benchmark for triangulating the comparison between MSNs and two other MRI-based networks measured in the same NSPN cohort: the structural covariance network (SCN) based on inter-regional correlations of a single feature (cortical thickness) measured across all 296 participants, and the sample mean DWI network based on tractographic reconstruction of white matter connections between cortical areas (weighted by mean diffusivity) in each participant (see **Methods**). To compare MSN, DWI and SCN results, we constructed a series of sparsely connected graphs (with connection density ranging from 0.5% to 5% in 0.5% increments) that represented the most strongly connected edges in each of the MRI networks and calculated the percentage of edges that linked areas in the same cytoarchitectonic class. For the most sparsely connected MSN (0.5% density), more than 90% of edges connected areas in the same class and this declined monotonically as a function of increasing density so that about 60% of edges connected areas in the same class in the MSN at 5% density. The SCN and DWI networks demonstrated similar trends but the percentage of intra-class connectivity was consistently lower for both these networks than for the MSN across all connection densities (Friedman χ^2^(2) = 40, *P* < 0.001) (**Figure 3B**). Only about 60% of edges in the DWI network connected areas of the same cytoarchitectonic class, even at the sparsest connection density, which was reflected in the relative failure of the DWI tractography to recover long distance connections between bilaterally symmetric areas belonging to the same class (**Figure 3C**). These comparative analyses demonstrate that morphometric similarity provided a closer approximation to the histological similarity between two cortical regions than analysis of either cortical thickness covariance or DWI-measures of white matter connectivity.

### Morphometric Similarity and Cortical Gene Co-Expression

We tested the second biological hypothesis, that MSN edges link areas with high levels of gene co-expression, using two gene sets: i) the approximately complete human genome (20,737 genes) and ii) a much smaller subset of 19 HSE (human supragranular enriched) genes that are known to be specifically expressed in the supragranular layers (cortical lamina II and III) of human cortex (Zeng et al., 2012) that are characteristic of the cytoarchitectonic classes (1-3) with higher nodal similarity scores (**Figure S5A**). We mapped the whole genome transcriptional data on 6 adult human post-mortem brains (Hawrylycz et al., 2012) into the same parcellation scheme that was used to define the 308 nodes of the MSNs (Whitaker and Vértes et al., 2016; Vértes et al., 2016) (**Methods**). Then we could estimate the inter-regional co-expression (Pearson correlation) of gene transcriptional profiles for each possible pair of nodes in the same anatomical frame of reference as the MSNs.

There was a significant positive correlation between the edge weights of the sample mean MSN and inter-regional co-expression of the whole-genome (*r* = 0.33, *P* < 0.001), meaning that cortical areas with high morphometric similarity also tended to have high transcriptional similarity. This correlation was attenuated, but remained statistically significant (*r* = 0.19, *P* < 0.001) after accounting for shared distance effects on inter-regional morphometric and transcriptomic similarity (see **Methods**).

To prioritise genes by the strength of their contribution to the observed association between morphometric similarity and whole genome co-expression, we used a leave-one-out procedure whereby the correlation between MSN edge weights and gene co-expression was iteratively re-estimated after systematically removing each one of the genes in turn. This algorithm allowed us to rank all 20,737 genes in terms of the difference their exclusion from the analysis made to the estimated correlation between morphometric similarity and gene co-expression (see **Supplemental Information** for the list of genes and their rankings). Gene ontology (GO) enrichment analysis of this gene list revealed that high-ranking genes – which made a stronger contribution to the association between morphometric and transcriptomic similarity - were enriched for annotations related to neuronal structure and signalling (**Figure S5C**). The HSE gene list of interest *a priori* was also high-ranking, with a median rank that was within the top decile of all genes (median HSE gene rank = 1,889/20,737), and significantly greater than the median ranks of 10,000 random gene sets of equal size (n=19, *P* < 0.0001). Moreover, nodal similarity of the sample mean MSN was positively correlated with regional gene expression (*r* = 0.34, *P*_*bootstrap*_ = 0.0042) and regional co-expression (*r* = 0.48, *P*_*bootstrap*_ = 0.01) of the HSE gene set (**Figure S5B**); and HSE gene expression demonstrated the same distribution across cyotarchitectonic areas as nodal morphometric similarity (**Figure S5A**). Specifically, expression of HSE genes was greater in cytoarchitectonic areas 1-3 where the hubs of the morphometric similarity networks were also concentrated. In a complementary analysis of phenotype enrichment in mammalian gene knockout models (Smith and Eppig, 2012) using Enrichr (Chen et al., 2013; Kuleshov et al., 2016), we found that disruption in animal models of high-ranking genes (i.e., those with positive leave-one-out scores) was significantly associated with abnormal synaptic transmission (*P* < 0.05, FDR corrected). Collectively, these results indicate that MSN topology is aligned with spatial expression patterns of neuronally-expressed genes that are enriched within human cortical layers mediating cortio-cortical connectivity, and genes that are critical for normal neuronal functions.

### Morphometric similarity (MRI) compared to tract tracing connectivity in the macaque

To assess the generalisability of MSN methodology to non-human primate MRI datasets, and to test the third biological hypothesis, that MSN edges indicate axonal connectivity between cortical areas, we analysed a publically-available collection of MRI data on a cohort of 31 juvenile rhesus macaque monkeys (16 females, overall age range = 0.9-3.0 years) (Young et al., 2017). We constructed MSNs for each monkey as described previously for each human participant (**Figure 1**), using 8 morphometric features available from the T1-weighted, T2-weighted and DWI data on each animal. We defined 91 nodes of the macaque cortex using a histologically-defined and anatomical landmark-based parcellation (Markov et al., 2014) that has been previously used to report retrograde axonal tract-tracing experiments on 29 of the 91 nodes (Markov et al., 2012; Markov et al., 2014) (see **Figure 4B**).

**Figure 4.**
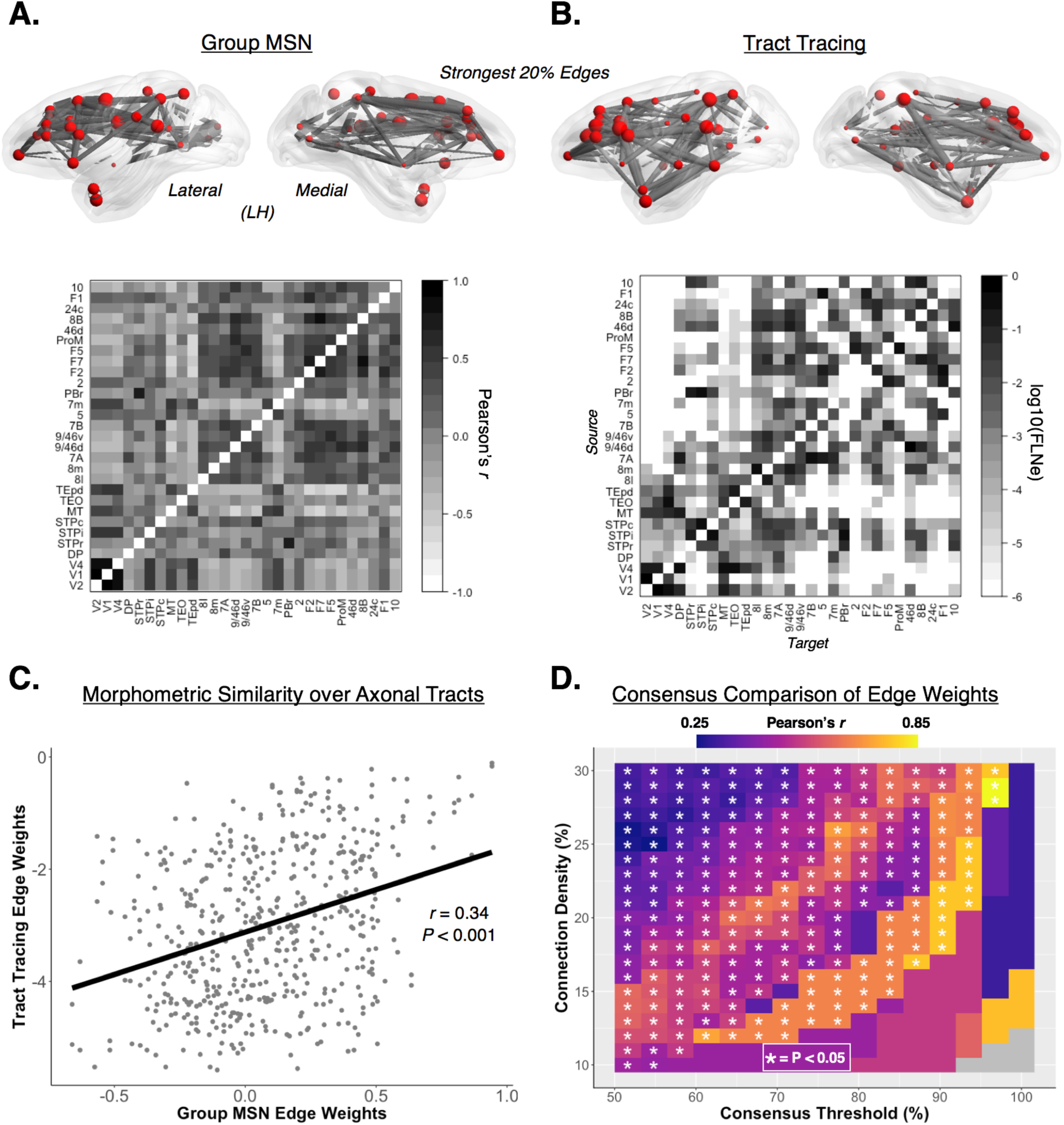
Comparison of morphometric similarity to axonal tract tracing in the macaque. (Top) The left hemisphere edges of the group average multimodal macaque MSN (**A**) and axonal tract tracing network (**B**), each thresholded at 20% connection density. Nodes in both networks are sized according to degree – calculated as the average nodal degree across MSNs (at 66% connection density) and, due to the effects of directionality, averaged nodal degree across both efferent and afferent connections in the tract tracing network. (Middle) The 29 × 29 group average multimodal macaque MSN (**A**) and the connections of the tract tracing matrix (**B**). The 29 × 29 tract tracing connectivity matrix is based on retrograde injections in 29 regions of the macaque cerebral cortex (Markov et al., 2012), and is 66% dense. Connection weights are based on the extrinsic fraction of labelled neurons (FLNe), and are plotted on a base 10 logarithmic scale. Diagonals in both networks are whited-out. **C)** For the overlapping edges in the two matrices in b), there was a significant positive correlation between the edges of the group macaque MSN and the edge weights of the tract tracing network (*r* = 0.34, *P* < 0.001). **D)** The correspondence between the edge weights of the group MSN and those of the tract tracing network. The group MSN was masked using a consensus approach, which incorporated the most common edges of the individual MSNs at varying connection densities (10-30%). At each connection density and consensus threshold (determined by the proportion of subjects at a connection density with common supra-thresholded edges) the group MSN and tract tracing network were masked and the edge weights were correlated. Generally, we observed a positive relationship in connectivity weights across MSN connection densities and consensus thresholds (median *r* = 0.58, range = 0.27-0.82), with the highest correlations found using the strongest and most consistent edges in the individual MSNs. Collectively, these results not only suggest a relationship between morphometric similarity (derived *in vivo*) and axonal tract weights (derived *ex vivo*) at the group level, but also reveal a possible “core” set of associations (measured in individual MSNs) which closely approximate physical anatomical connectivity.

Macaque MSNs demonstrated qualitatively the same suite of complex topological properties as the human MSNs from both the NSPN and NIH cohorts (**Figure 4A, Figure S3**). There was a significant positive correlation between the edge weights of the sample mean macaque MSN and the edge weights of the tract-tracing network (Pearson’s *r* = 0.34, *P* < 0.001) (**Figure 4B/C**). To test if this relationship varied as a function of MSN edge strength and consistency across individuals, the correlation between MSN and tract-tracing edge weights was estimated across a range of MSN connection densities (10-30%), and edge consistencies across individuals (50-100%). This analysis revealed that edge weights of the individual MSNs, consistently evident in the more sparsely connected graphs, were strongly correlated with the anatomical connectivity weights derived from axonal tract tracing data (Pearson’s *r* = 0.36-0.90, median *r* = 0.58, 73% of correlations *P* < 0.05, 34% of correlations Bonferroni-corrected *P* < 0.05) (**Figure 4D**). Taken together, these findings indicate that the morphometric similarity of two cortical regions is directly related to the strength of monosynaptic axonal connectivity between them.

### Predicting individual differences in human IQ from differences in nodal degree of morphometric similarity networks

Having established the technical feasibility, robustness, and biological validity of morphometric similarity mapping, we leveraged the ability to MSNs to represent whole brain anatomical networks in a single human to investigate relationships between inter-individual differences in brain network topology and inter-individual differences in cognitive and behavioural traits.

We focused on general intelligence (IQ) as the cognitive trait of interest given the broad relevance of IQ for adaptive function (Davies et al., 2016; Hagenaars et al., 2016) and the wealth of prior research into biological substrates for IQ (Crossley et al., 2013; Dehaene and Changeux, 2011; van den Heuvel et al., 2009). We predicted that IQ should be positively associated with integrative topological features that promote efficient information transfer across the whole network. High degree hub nodes are crucial to the global efficiency of the connectome and preferentially impacted by clinical brain disorders associated with cognitive impairment (Crossley et al., 2014). On this basis, we predicted specifically that individual differences in verbal and non-verbal IQ should be related to individual differences in nodal degree.

We used the multivariate technique of partial least squares (PLS) regression to find the optimal low-dimensional relationship between a set of predictor variables and response variables. In our case, the (292 × 308) predictor variable matrix comprised measurements of degree (calculated at 10% connection density) at each of 308 nodes in each of 292 participants in the NSPN cohort; the (292 × 2) response variable matrix comprised *t*-scores on the verbal (vocabulary) and non-verbal (matrix reasoning) scales of the Wechsler Abbreviated Scale of Intelligence (WASI; Wechsler, 1999), standardised for age and gender effects relative to a representative population, and measured in the same 292 participants (mean total IQ = 111, SD = 12, range = 76-137). Each of the predictor variables (MSN degree) was regressed on the potentially confounding effects of age, gender, and age × gender interaction, before the residuals were used in the PLS analysis.

The first two components of the PLS explained about 40% of the variance in IQ, and this goodness of fit was statistically significant by a non-parametric resampling procedure (*P* = 0.03) (Vértes et al., 2016). These results maintained statistical significance (*P* < 0.05) when the analysis was repeated for nodal degree calculated across a range of MSN connection densities (10-25%). We focus our attention on the first two PLS components (PLS1 and PLS2), which consistently explained about 25% and 15% of the variance in IQ, respectively (**Figure 5**).

**Figure 5.**
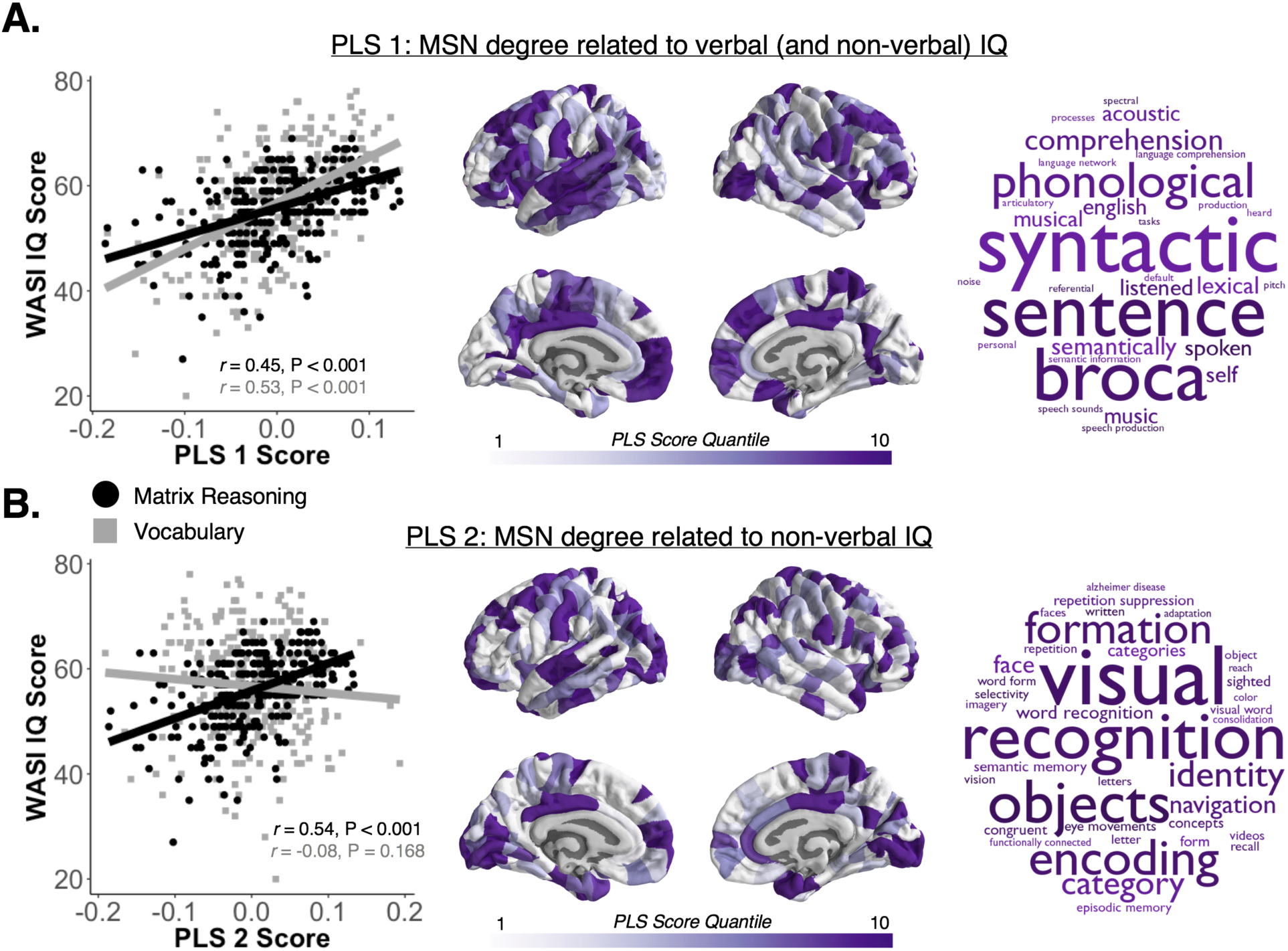
Nodal degree of morphometric similarity networks is highly predictive of individual differences in intelligence. The first two components (PLS1, PLS2) of a partial least squares regression using individual MSN degree (at 10% connection density) explained about 40% of the variance in vocabulary and matrix reasoning subscales of WASI IQ scores in 292 people. **A)** The first PLS component (PLS1) was correlated with both vocabulary and matrix reasoning (Left) and with the degree or hubness of nodes in left-lateralised temporal and bilateral frontal cortical areas (Centre), related to language functioning (Right). **B)** PLS2 was correlated specifically with matrix reasoning and degree or hubness of nodes in bilateral primary sensory cortical areas (Centre), specialised for visual and sensorimotor processing.

The first PLS component was significantly positively correlated with both IQ subscales - vocabulary (*r* = 0.53, *P* < 0.001) and matrix reasoning (*r* = 0.45, *P* < 0.001), as well as full-scale IQ (*r* = 0.61, *P* < 0.001) (**Figure 5B**). The second PLS component was only significantly positively correlated with matrix reasoning (*r* = 0.54, *P* < 0.001) and full-scale IQ (*r* = 0.21, *P* < 0.001), but not vocabulary (*r* = 0.08, *P* = 0.168) (**Figure 5B**). We ranked the 308 nodes of the individual MSNs according to their bootstrap standardised weight on each PLS component (Vértes et al., 2016). This analysis revealed that the high degree hubs that loaded strongly on PLS1, and were positively correlated with higher verbal and non-verbal IQ, were located predominantly in left frontal and temporal cortical areas; whereas the high degree hubs that loaded strongly on PLS2, and were positively correlated with non-verbal IQ, were located predominantly in bilateral occipital and frontal cortex. We used Neurosynth, a tool for meta-analysis of the large primary literature on task-related fMRI (Yarkoni et al., 2011), to identify which cognitive functions were co-localised with the cortical nodes strongly weighted on PLS1 and PLS2. As expected, high degree hubs located in frontal and temporal cortex and predictive of verbal and non-verbal IQ were enriched for language-related functions; whereas high degree hubs located in occipital cortex and predictive of non-verbal IQ were enriched for visual and memory functions (**Figure 5**).

## Discussion

We have shown how multimodal MRI measurements of human and non-human primate cortex can be used to estimate the morphometric similarity between cortical areas and the topological properties of the anatomical connectome of a single brain. This robust and replicable new method of brain structural network analysis allowed us to test (and affirm) three key biological hypotheses about the organisation of individual mammalian cortical networks. As theoretically predicted, we found evidence that cortical areas connected by an edge in morphometric similarity networks were cytoarchitectonically similar and axonally connected to each other, and had high levels of co-expression of genes specialised for neuronal functions. These results substantiated the biological validity of MSNs, compared to other MRI or DWI-based estimates of the human connectome, and motivated us to test (and affirm) a fourth hypothesis: that individual differences in IQ are related to individual differences in the hubness or nodal degree of cortical nodes in human brain anatomical networks.

Like most spatially embedded real-life networks, including other brain networks, morphometric similarity networks had a complex topology (Barabási, 2016; Fornito et al., 2016). MSNs were binary graphs with a small-world combination of high clustering and short path length; some high degree hub nodes with many connections to the rest of the network; and a community structure comprising modules and a rich club. This suite of topological properties was robust to variation in species (human and macaque), human volunteer samples (NSPN and NIH cohorts), static magnetic field strength for MRI (1.5T and 3T), number of morphometric features measured by MRI at each regional node (5, 8 or 10 MRI-derived parameters per region), as well as pre-processing steps like cortical parcellation.

This robust and replicable network phenotype or connectome, derived from morphometric similarity mapping, is qualitatively similar to connectomes previously described using comparable graph theoretical metrics in many other neuroimaging and neuroscience datasets. A well-rehearsed interpretation of the complex topology of connectome organisation is in terms of its supposed advantages for sensory, motor or cognitive function. Some topological features, such as clusters and modules, will favour segregated processing of specific channels of information, whereas other features, such as hubs and a rich club, will favour integrated processing of all information (Bullmore and Sporns, 2012; Sporns et al., 2004). This influential hypothesis, linking the topology of the human connectome to the psychological capacities of the brain, has some experimental support. The evidence is strongest for the link between modular or clustered topologies and specialised psychological or information processing functions (Fodor, 1983). At all scales of connectomics, from micro-scale analysis of *C. elegans* or *Drosophila* to macro-scale analysis of human MRI data (Meunier et al., 2010; Schröter et al., 2017), there is evidence for topological modules of spatially co-localised (neuronal or areal) nodes with specialised functions. The evidence is not yet so strong for the link between integrative topological features – such as hubs and a rich club – and global or generalised cognitive functions (we return to this point later).

### Morphometric similarity and anatomical similarity

By aligning individual morphometric similarity networks with the classical cytoarchitectonic atlas of von Economo and Koskinas, we demonstrated a close alignment between MSN topology and this histological classification of cortical areas. Morphometric similarity, measured by MRI, was greater between regional nodes that were histologically similar in the sense of belonging to the same cytoarchitectonic class. This meant that sparse MSNs, representing only a small percentage of the highest morphometric similarity statistics, were dominated by intra-class edges between regions. Correspondence between morphometric similarity and cytoarchitectonic similarity is supportive of the biological validity of the constituent MRI measurements. There is also growing evidence that cytoarchitectonic similarity predicts axonal connectivity between cortical areas, with greater probability of axonal connectivity between histologically similar areas (Goulas et al., 2017; Goulas et al., 2016). Thus, we reasoned, alignment of network edges with cytoarchitectonic classes could provide a triangulation point to compare MSNs to other MRI-based methods of human connectome mapping. Since histologically similar nodes are more likely to be axonally connected, then any map of anatomical connectivity derived from MRI should be dominated by intra-class edges.

We compared MSNs to two other MRI-based anatomical networks estimated from the same sample – a single structural covariance network and a set of individual diffusion tractography networks. All three networks had qualitatively and quantitatively similar complex topology, but they were not identical. In relation to the benchmark of cytoarchitectonic classification, all networks were dominated by a high percentage of intra-class edges when graphs were thresholded sparsely to include only the strongest connections between regions. However, across all connection densities considered, the percentage of intra-class edges was greater for MSNs than for the SCN or DWI networks. This indicates that edges in MSNs are more representative of histologically similar pairs of regions, which are more likely to be axonally connected to each other, than edges in the SCN or DWI networks. One reason for the relatively poor performance of DWI networks in aligning to cytoarchitectonic standards seems likely to be the known difficulties in reconstructing long distance axonal connections by tractography analysis of DWI data. This would explain why DWI networks, compared to both MSNs and the SCN, relatively lacked inter-hemispheric intra-class edges between homologous cortical areas in the same cytoarchitectonic class.

Although cytoarchitectonic classification is a well-established and traditional way of assessing histological similarity between cortical areas, more generally we can assess inter-areal similarity in terms of any locally (spatially) expressed cellular or genomic phenotype. Spatial patterns of gene expression in the mammalian cortex are intimately tied to regional differences in cortical layering and cell-composition (Bernard et al., 2012; Hawrylycz et al., 2012). Transcriptomic similarity or gene co-expression was greater between regions of the mouse brain that were known to be axonally connected by analysis of anterograde tract-tracing data (Fulcher and Fornito, 2016). A functionally specialised set of so-called HSE genes, that are over-expressed specifically in supragranular layers of human association cortex (Zeng et al., 2012), and known to be important for formation of long distance inter-areal axonal connectivity (Hawrylycz et al., 2012), were more strongly co-expressed by functionally connected brain regions (Krienen et al., 2016).

In this context, we predicted that morphometrically similar regions should have high levels of gene co-expression in general, as well as high levels of HSE gene co-expression in particular. Whole genome analysis confirmed that co-expression was positively correlated with morphometric similarity and the genes that contributed most strongly to the overall association between transcriptional and morphometric similarity were specialised for neuronal functions. HSE genes were most strongly expressed in cytoarchitectonic classes 1-3 and HSE gene co-expression was positively correlated with morphometric similiarity and degree or hubness of MSN nodes.

To this point, we have highlighted results that show morphometric similarity is strongly associated with cytoarchitectonic and genomic measures of histological similarity between cortical areas. To the extent that histological (cytoarchitectonic or transcriptional) similarity is coupled to axonal connectivity between cortical areas (Fulcher and Fornito, 2016; Goulas et al., 2017; Goulas et al., 2016), we can therefore expect morphometric similarity measured by MRI to be at least an approximate marker of axonal connectivity. However, to verify this important interpretation more directly, we generalised the MSN approach to analysis of whole brain connectomes in the macaque monkey.

We observed a strong positive relationship between the edge weights of the macaque MSN and the edge weights of the tract-tracing network, especially for the most consistently and strongly-weighted edges in the individual macaque MSNs. The strength of association between tract-tracing and morphometric similarity networks was comparable in magnitude to previous reports of correspondence between tract-tracing and DWI-based networks in the macaque (Donahue et al., 2016; van den Heuvel et al., 2015). We note that the macaque MRI data were collected at 3T and provided only 8 morphometric variables per cortical region. It is predictable from the human MRI datasets we have analysed that MSN metrics (and their alignment to tract-tracing data) could be more precisely measured in future macaque MRI experiments at higher field strength or using multi-parameter MRI sequences to sample cortical micro-structure more comprehensively.

In short, the results of multiple experiments convergently supported all three hypotheses linking the topology of morphometric similarity networks to the histological similarity and the axonal connectivity between cortical areas.

### Morphometric similarity networks and intelligence

The availability of a new method for mapping the anatomical connectome of a single human is likely to be helpful in understanding how its network topology relates to the cognitive or psychological functions of the brain. As noted earlier, it is particulalrly important to understand more clearly how integrative elements of connectome topology – like hubs and a rich club – might be linked to cognitive processing.

There is a body of theoretical and experimental work in support of the idea that higher order, more effortful conscious processing depends on a global workspace architecture that coordinates neuronal activity across anatomically distributed areas of cortex (Baars, 1997; Dehaene and Changeux, 2011). Conceptually related work has highlighted the importance of a multiple demand network of association cortical areas for fluid intelligence (Duncan, 2010). In the language of graph theory, this is compatible with the prediction that topologically integrative features of the connectome – which “break modularity” (Dehaene and Naccache, 2001) - should be important for intelligent cognitive function; and there is already some evidence in support of this prediction. For example, it has been shown that higher IQ is negatively correlated with the characteristic path length of fMRI and DWI networks (Li et al., 2009; van den Heuvel et al., 2009); that performance of a cognitively demanding working memory task was associated with greater topological efficiency (shorter path length) of MEG networks (Kitzbichler et al., 2011); and that the rich club of interconnected hubs in a meta-analysis of task-related fMRI data was co-activated by executive tasks demanding both cognition and action (Crossley et al., 2013).

On this basis, we predicted that individual differences in IQ should be related to variability in the hubness or degree of individual nodes in individual MSNs. We applied the multivariate, dimension-reducing method of partial least squares (PLS) to test the strength of association between the verbal and non-verbal IQ of 292 healthy participants, on one hand, and the degree of 308 cortical nodes in each of 292 individual MSNs, on the other hand. Remarkably, the first two PLS components collectively accounted for approximately 40% of the total variance in IQ. The first PLS component defined a set of cortical areas, functionally specialised for language and located in left frontal and temporal cortex, where higher degree was strongly predictive of higher verbal and non-verbal IQ; the second PLS component defined a distinct set of areas, functionally specialised for vision and memory, where higher degree was specifically predictive of higher non-verbal IQ.

These results directly support our fourth hypothesis and they provide some of the strongest evidence yet available in support of the more general hypothesis that topologically integrative features of the human brain connectome are important for “higher order” cognitive functions (Deary et al., 2010; Saggar et al., 2015). Indeed the strength of association between IQ and MSN nodal degree is large compared to many prior studies reporting an association between IQ and other structural MRI phenotypes (Reiss et al., 1996; Ritchie et al., 2015; Toga and Thompson, 2005). We predict on this basis that morphometric similarity mapping could provide a powerful technical platform for measuring the anatomical connectome in vivo and for understanding how the cognitive functions of the human brain are related to its topologically complex connectome.

## Author Contributions

Conceptualization, JS, AR, and ETB; Methodology, JS, FV, MS, AR, ETB; Validation, JS, AM, DL, and AR; Investigation, JS, FV, MS, RR-G, KJW, and PEV; Resources, AM, DL, PF, RJD, PBJ, IMG, AR, and ETB; Data Curation, JS, FV, RR-G, KJW, PEV, PKR, and LC; Writing – Original Draft, JS, FV, AR, ETB; Writing – Review & Editing, all authors; Project Administration, PF, RJD, PBJ, IMG, AR, ETB.

## Acknowledgments

We would like to thank the participants in the NSPN and NIH studies of human brain development. We also thank Claus Hilgetag, Alexandros Goulas, Konrad Wagstyl, Lisa Ronan, and James Rowe for helpful comments.

## Methods

We first summarise methods relevant to the principal human (NSPN) dataset and its analysis; then methods relevant to the secondary human (NIH) dataset; and finally methods relevant to the macaque monkey dataset.

### Study design and MRI data acquisition – primary NSPN cohort

Subjects were recruited as a part of the NeuroScience in Psychiatry Network (NSPN) study of normative adolescent development. A subgroup of 300 adolescents and young adults aged 14-24 years was stratified into 5 temporally contiguous strata: 14-15 years inclusive, 16-17 years, 18-19 years, 20-21 years, and 22-24 years (N=60 subjects per bin, 30 males and 30 females). Participants were excluded if they were currently being treated for a psychiatric disorder or for drug or alcohol dependence; had a current or past history of neurological disorders or trauma including epilepsy, or head injury causing loss of consciousness; had a learning disability requiring specialist educational support and/or medical treatment; or had a safety contraindication for MRI. Participants provided informed written consent for each aspect of the study and parental consent was obtained for those aged 14-15 years. The study was ethically approved by the National Research Ethics Service and was conducted in accordance with NHS research governance standards.

The anatomical MRI data were acquired using the multi-parametric mapping (MPM) sequence (Weiskopf et al., 2013) implemented on three identical 3T whole-body MRI systems (Magnetom TIM Trio, Siemens Healthcare, Erlangen, Germany; VB17 software version), two located in Cambridge and one in London, operating with the standard 32-channel radio-frequency (RF) receive head coil and RF body coil for transmission. Between-site reliability of all MRI procedures was satisfactorily assessed by a pilot study of 5 healthy volunteers scanned at each site (Weiskopf et al., 2013). The between-site bias was less than 3%, and the between-site coefficient of variation was less than 8%, for both the longitudinal relaxation rate (R1 = 1/T1) and MT parameters (Weiskopf et al., 2013). R1 and MT were quantified in Matlab (The MathWorks Inc., Natick, MA, USA) using SPM8 (www.fil.ion.ucl.ac.uk/spm) and custom tools (Weiskopf et al., 2013; Whitaker and Vértes et al. 2016).

Diffusion weighted imaging (DWI) data were collected in the same scanning session as the MPM data. A High-Angular Resolution Diffusion-Weighted Image (HARDI) was acquired using a single-shot echo planar imaging sequence consisting of 63 gradient directions with a b-value = 1000 mm/s2 along with 5 unweighted B0 images. This protocol used 70 consecutive axial slices of thickness 2 mm (FOV=192 × 192 mm, TE=90 ms, TR= 8700 ms) resulting in a voxel size of 2.0 mm isotropic.

### Human MRI data pre-processing

We used FreeSurfer v5.3.0 software for the data pre-processing pipeline (http://surfer.nrm.mgh.harvard.edu). Briefly, the cortical surface for each participant was reconstructed from their R1 image by the following steps: skull stripping (Segonne et al., 2004), segmentation of cortical grey and white matter (Dale et al., 1999), separation of the two hemispheres and subcortical structures (Dale et al., 1999; Fischl et al., 2002; Fischl et al., 2004); and finally construction of smooth representation of the grey/white interface and the pial surface (Fischl et al., 1999). The DWI volumes were aligned to the R1 image for each subject. After quality control, 3 participants had to be excluded from the analyses due to movement artefacts which prevented accurate surface reconstructions, and 1 due to errors in their DWI volume reconstruction, leaving N=296 (148 males and 148 females) for the final cohort used for the imaging analyses in this paper.

### Human MRI cortical parcellation

To define the set of nodes, the 68 cortical regions in the Desikan-Killiany atlas (Desikan et al., 2006) were sub-parcellated into 308 spatially contiguous regions, of approximately equal size (∼5 cm^2^), using a backtracking algorithm as described previously (Romero-Garcia et al., 2012). This parcellation was generated once on the surface of the FreeSurfer standard anatomical template (fsaverage), and subsequently transformed to each individual subject’s surface. Each subject’s surface parcellation was then interpolated and expanded to their respective R1, MT, and B0 (DWI) volumes.

### Estimation of regional morphometric features

A feature matrix consisting of 10 morphometric features measured at each of 308 brain regions was estimated from the combined MPM and DWI data available for each subject (**Figure 1A**). Surface- and volume-based features were extracted using the respective version of the regional parcellation. For the surface-based features, regional values were estimated for cortical thickness (CT), surface area (SA), intrinsic curvature (IC), mean curvature (MC), curvature index (CI), and folding index (FI). For the volume-based features, regional values were estimated for the diffusion metrics (fractional anisotropy – FA, and mean diffusivity – MD) as well as grey matter volume (GM) and magnetization transfer (MT). The regional MT values were estimated at 70% cortical depth (Whitaker and Vértes et al., 2016).

### Estimation of morphometric similarity and morphometric similarity networks

Each of the MRI feature vectors in each region of each individual image were normalised (z-scored) and then the Pearson product-moment correlation coefficient (*r*) was estimated for each possible regional pair of MRI feature vectors (see **Figure 1B/C**). These pair-wise measures of morphometric similarity were compiled to form a morphometric similarity matrix which was thresholded to construct binary graphs of arbitrary connection density, also known as morphometric similarity networks (MSNs). Explicitly, MSNs were thresholded such that inter-regional correlations less than the threshold value were set to 0, and supra-threshold edges were set to 1, in the corresponding elements of the individual MSNs. To create a group level MSN, we averaged the individual morphometric similarity matrices and then thresholded the mean similarity matrix.

### Structural covariance network construction

Cortical thickness (CT) of each of the 308 regions in the 296 subjects was used as the morphometric feature for constructing the structural covariance network (SCN), as in Whitaker and Vértes et al. (2016). This entailed estimating the set of correlations between cortical thickness of each possible pair of regions, over all participants, resulting in a single inter-regional CT correlation matrix which was thresholded to generate a binary graph representation of the sample mean structural covariance network.

### DWI network construction

The entire DWI image analysis pipeline was performed in AFNI, a freely available MRI and DWI analysis software suite (Cox, 1996). For each subject, DWI image volumes were de-obliqued and co-registered to the B0 volume to account for head movement. The 6 principal direction tensors were then estimated from the DWI image volume using the 3dDWItoDT command. Probabilistic tractography for the 308 brain regions was performed using the 3dTrackID command, along with tensor uncertainty estimates from 3dDWUncert to increase robustness (Taylor et al., 2012; Taylor and Saad, 2013). A connectivity matrix was estimated for each of the 296 subjects, where each element represents the average mean diffusivity (MD) along the axonal tracts connecting two regions. We chose the measure for estimating tract weights that maximised the proportion of intra-class edges defined by a prior cytoarchitectonic classification (**Figure 3).**

### Graph theoretic analyses

For each of the individual MSNs (and comparable SCN and DWI networks), a series of graphs was constructed and analysed over a range of connection densities (10-40%, 5% intervals). Summary statistics for the MSNs, as well as for random networks with the same number of nodes and edges, are reported using the 7 arbitrary thresholds.

The following graph metrics were calculated in Matlab using the Brain Connectivity Toolbox, a freely available graph analysis software package (Rubinov and Sporns, 2010), as well as in R (R Development Core Team, 2008, http://www.r-project.org/) using the igraph package (Csardi and Nepusz, 2006). <u>Graph visualisation was performed using custom code written in R and Python (Python Software Foundation,</u> http://www.python.org/), as well as using BrainNet Viewer (Xia et al., 2013) (http://www.nitrc.org/projects/bnv/).

The degree, *k*, of a graph describes the total number of edges of each node. The clustering coefficient, *C*, of node *i* is the ratio of the number of *i*’s neighbours that are connected to each other with a single edge. As such, the clustering coefficient across an entire graph is the average of the clustering coefficients over all nodes, defined as:

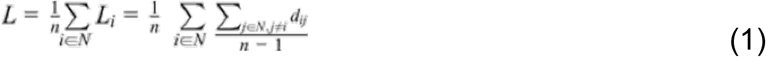

where *k*_*i*_ are the neighbours of node *i* and *k*_*i*_(*k*_*i*_ *– 1*) is the number of possible edges between *k*_*i*_ (Watts and Strogatz, 1998). The characteristic path length represents the mean of shortest paths between all pairs of nodes in a network (Watts and Strogatz, 1998), defined as:

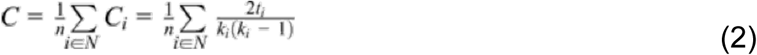

where *L*_*i*_ is the mean distance between node *i* and every other node. Global efficiency is the inverse of the characteristic path length. Modularity (*Q*) is a measure of network segregation, by which nodes are subdivided into communities, or modules, to maximise the number of within-module connections while minimising the number of between-module connections (Girvan and Newman, 2002; Newman, 2004a, b). It is defined as:

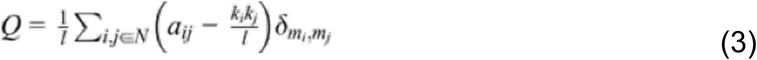

where *m*_*i*_ and *m*_*j*_ are the modules containing nodes *i* and *j*, respectively. If *m*_*i*_ = *m*_*j*_,(i.e. nodes *i* and *j* are members of the same module) *then* δ*m*_*i*_,*m*_*j*_ = 1, but if *m*_*i*_ ≠ *m*_*j*_, (i.e. nodes *i* and *j* are members of different modules), then δ*m*_*i*_,*m*_*j*_ = 0. The small-world coefficient of a network (Humphries and Gurney, 2008) is defined as:

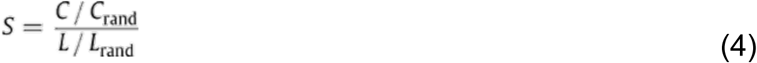

where (*C/C*_*rand*_) is the ratio of the average clustering coefficient between the empirical network and average values from a set of corresponding randomised networks with preserved degree distribution, and (*L/L*_*rand*_) is the ratio of characteristic path lengths of the empirical network and the set of random networks. In this study, 1 random network per participant was generated. As such, small-world networks (generally *S* > 1) have high clustering and similar or shorter path lengths compared to random networks.

The rich club coefficient is a property of complex networks and measures the amount of inter-connectedness between hubs of a network (Colizza et al., 2006; van den Heuvel and Sporns, 2011), calculated at varying degree cut-offs. The normalised rich club coefficient (*Φ*) is the ratio of the rich club coefficient of the empirical network (i.e., the MSNs) relative to that of a random network. Thus the nodes greater than or equal to the degree cut-off at which *Φ* > 1 denotes membership to the rich club. We present the rich club results of the MSNs at a 10% connection density.

### Digitisation of a cytoarchitectonic atlas

To test the correspondence between morphometric similarity and cytoarchitecture, we used an independent modular decomposition depicting the five typical laminar patterns of the cortex as proposed by von Economo and Koskinas (1925). We manually assigned nodes in our N=308 parcellation to one of the five cortical classes (Solari and Stoner, 2011), and, in addition, the insula and cingulate cortex were partitioned into separate classes to reflect their distinct cytoarchitectonic profiles, thus producing seven distinct modules (Vertes et al., 2016; Whitaker et al., 2016).

### Brain gene co-expression estimation

We used the freely available Allen Institute of Brain Sciences (AIBS) transciptomic dataset (http://human.brain-map.org/) to estimate gene expression for each region and gene co-expression between each pair of regions. This dataset comprises post-mortem samples collected from 6 adult male donors (H0351.1009, H0351.1016, H0351.1015, H0351.2002, H0351.1012, H0351.2001; 3 Caucasian, 2 African-American, 1 Hispanic; mean age = 42.5 years). For a detailed description on the methods of whole genome microarray analysis at multiple regional locations, see the AIBS technical white paper (http://human.brain-map.org).

To account for the redundancy of the cRNA hybridization probes, which contained expression levels for overlapping genes, expression values for the same gene were averaged across probes. Probes with unmatched genes were excluded, leaving 20,737 genes from 3,702 samples. Because of the symmetric gene expression values between hemispheres (Pletikos et al., 2014), the AIBS dataset only contains data from both hemispheres for two subjects. Thus, because the right hemisphere was under-sampled, we performed all analyses on the left hemisphere (N = 152 regions) by reflecting the right hemisphere samples. We then mapped the gene expression values of each subject to the fsaverage (MNI305) volumetric template space (assigning samples to the nearest centroid of the left hemisphere (N=152 regions) of our parcellation) using the individual AIBS subjects’ T1-weighted volumes (Vertes et al., 2016; Whitaker et al., 2016).

The median regional expression was estimated for each gene across participants (N=6) and then each gene’s regional values were normalised (z-scored), resulting in a 152 × 20,737 matrix of the genome-wide expression data for the 152 regions of the left hemisphere. The 152 × 152 gene co-expression matrix (i.e., the upper-left quadrant of the group MSN) contained pairwise Pearson correlations, which were computed for each of the left hemisphere regions, representing the intra-hemispheric gene co-expression of two left hemisphere regions across the 20,737 genes. This network, along with the set of regional expression values, was used for comparison to the corresponding left hemisphere of the group MSN.

The Human Supragranular Enriched (HSE) gene set contains 19 genes that were found to be primarily expressed in the upper layers (II/III) of human cortex: *BEND5, C1QL2, CACNA1E, COL24A1, COL6A1, CRYM, KCNC3, KCNH4, LGALS1, MFGE8, NEFH, PRSS12, SCN3B, SCN4B, SNCG, SV2C, SYT2, TPBG* and *VAMP1* (Zeng et al., 2012). The inter-areal co-expression of HSE genes has been related to the emergence of cortico-cortical connectivity in humans (Krienen et al., 2016; Zeng et al., 2012). We therefore created a gene co-expression network using only the HSE genes.

Gene ontology enrichment analysis was performed using GOrilla (Eden et al., 2007; Eden et al., 2009) and visualised using REViGO (Supek et al., 2011), and, in addition, replicated using Enrichr (Chen et al., 2013; Kuleshov et al., 2016).

### Partial least squares analysis

We assessed the relationship between individual differences in IQ and individual differences in nodal degree of each of 308 regions in each of 292 individual MSNs using the multivariate method of partial least squares (PLS) regression, as in Whitaker and Vértes et al. (2016) and Vértes et al. (2016). This dimensionality reduction technique seeks to find the latent variables or PLS components which maximise the correlation between a set of collinear predictor variables and a set of response variables. Here, normalised scores on the vocabulary and matrix reasoning subscales of the Wechsler Abbreviated Scale of Intelligence (WASI) test (Wechsler, 1999) were used as response variables and degree centrality of each node in individual MSNs thresholded at arbitrary connection density were used as predictor variables. We used MSN degree so that only high strength edges were included in the analysis, and so that the robustness of this PLS method could be tested across a range of MSN densities. Four subjects were excluded due to unavailability of WASI IQ data (N=292). Bootstrapping (resampling with replacement of the 292 individual subjects) was used to estimate the error on the PLS weights for each node so that the nodes could be ranked based on their contribution to each PLS component (Vértes et al., 2016).

### Study design and MRI data acquisition – secondary NIH cohort

To test the reliability and replicability of morphometric similarity, we constructed MSNs in an independent cohort of human subjects, using different image analysis software for the processing pipeline, a different areal parcellation, and a more limited set of morphometric features (N=5). The sample consists of 124 typically developing subjects (49 females, mean age = 12.55, σ = 4.27, range = 5.59-25.13; 65 males, mean age = 13.40, σ = 4.52, range = 5.77-32.04) sampled from the National Institutes of Health (NIH) longitudinal study of normative brain development (Evans and Brain Development Cooperative, 2006; Giedd et al., 1999; Giedd et al., 2015). Briefly, subjects were scanned on a 1.5T GE Signa scanner (axial slice = 1.5 mm, TE = 5 ms, TR = 24 ms, flip angle = 45°, matrix = 256x256x124, fov = 24 cm) using a spoiled-gradient recalled echo (3D-SPGR) imaging sequence (Giedd et al., 1999). The T1-weighted scans were processed using the Montreal Neurological Institute’s CIVET pipeline (Ad-Dab’bagh et al., 2006). Each of the scans and surface reconstructions of the 124 subjects were subjected to and passed quality-control assessment by three independent raters. Due to the lack of MT, DWI, or T2-weighted imaging, only (grey matter) morphometric features derived from the T1-weighted scans were estimated (CT, SA, GM, MC, IC). GM values were estimated using the T1w volumes of each subject. Vertex-wise CT and SA values were estimated using the resultant pial surface reconstructions from CIVET, while MC and IC metrics of these surfaces were estimated using the freely available Caret software package (Van Essen et al., 2001). The down-sampling of theses surface meshes (∼80,000 vertices per mesh) into 360 regions was performed (Alexander-Bloch et al., 2013b), where the vertex-wise estimates of the features were averaged (for CT, MC, and IC) or summed (for SA) within a given region in the parcellation. The surface parcellation was projected to the volume for extraction of regional GM for each subject.

### Macaque monkey MRI data acquisition

We additionally constructed MSNs using an independent cohort of 31 healthy young rhesus macaques (13 females; mean age = 1.7 years, range = 0.9 to 3.0 years), whose MRI data were collected as part of the UNC-UW longitudinal study at the University of North Carolina and the University of Wisconsin (Young et al., 2017). Animals were included if they had T1-weighted, T2-weighted and DWI data available at more than one time point. Prior to scanning, each animal was anaesthetised following the protocol from Young et al. (2017). All animal procedures were conducted in compliance with the Institutional Animal Care and Use Committee (IACUC) and the National Institutes of Health Guide for the Care and Use of Laboratory Animals.

Briefly, T1-weighted (*TR* = 8.684 ms, *TE* = 3.652 ms, FOV = 140 × 140 mm, flip angle = 12°, matrix = 256 × 256, thickness = 0.8 mm, gap = -0.4 mm, voxel resolution = 0.55 × 0.55 × 0.8 mm^3^) and T2-weighted (*TR* = 2500 ms, *TE* = 87 ms, FOV = 154 × 154 mm, flip angle = 90°, matrix = 256 × 256, thickness = 0.6 mm, gap = 0 mm, voxel resolution = 0.6 × 0.6 × 0.6 mm^3^), and diffusion-weighted images (120 gradient directions, *TR* = 8000 ms, *TE* = 65.7 ms, FOV = 16.7 mm, matrix = 128 × 128, thickness = 2.6 mm, voxel resolution = 0.65 × 0.65 × 1.3 mm^3^) were acquired using a GE MR750 3T scanner with an 8-channel human brain array coil (Young et al., 2017).

Following the processing pipeline in, the same 5 morphometric features used to construct the NIH human MSNs were estimated for the macaque data using an automated analysis pipeline (Seidlitz et al., 2017), which combines tools from AFNI as well as from the freely available Advanced Normalization Tools (ANTs) software package (Avants et al., 2011) (<u>picsl.upenn.edu/software/ants/</u>). Additionally, we estimated MD and FA from the DWI scans, as well as the T1w/T2w ratio (after resampling the T2w scans to match the resolution of the T1w scans) (Glasser and Van Essen, 2011), generating a total of 8 regional morphometric features for each subject. For cross-species comparison of the network properties, and for comparison to retrograde viral tract tracing data, we used the 91-region left hemisphere cortical parcellation from Markov et al. (2014). The individual 91 × 91 region left hemisphere MSNs were averaged to create a group average macaque MSN.

### Macaque tract tracing data

Connectivity of the group MSN was evaluated against tract tracing connectivity data from Markov et al. (2012) (downloaded from core-nets.org). Whereas DWI tractography is an *in vivo* non-invasive indirect approximation of white matter connectivity, “gold standard” anatomical tract tracing is an invasive *ex vivo* method of measuring directed connectivity. In retrograde tracing, a tracer (typically a dye, molecule, or radioactively-tagged amino acid) is injected and physically travels along the axonal projection from its termination site (i.e. site of injection) to the soma from which the axon originates. The measure of connectivity is reliant upon the labelling of neurons in the areas of interest. Markov et al. (2012) generated an index of connectivity called the extrinsic fraction of labelled neurons (FLNe). FLNe was calculated as the number of labelled neurons at the target site that exist above and beyond the fraction of labelled neurons of that site relative to the entire brain and labelled neurons intrinsic to that site (Markov et al., 2012). Thus, this retrograde tract tracing dataset is a weighted and directed 29 × 91 matrix (i.e. 29 injection points), where the 29 × 29 subgraph of the matrix contains all possible connections between those injection sites.

The distribution of edge weights from the tract tracing network follows a logarithmic distribution, thus we transformed the edge weights by the base 10 logarithm function for our analyses. We used the 29 × 29 subgraph of the total 29 × 91 connectivity matrix (**Figure 4B**). For comparison with the tract tracing data, the 91 × 91 macaque MSNs were matched to the same 29 × 29 dimensions.

